# Antitumor Activities and Cellular Changes Induced by TrkB Inhibition in Medulloblastoma

**DOI:** 10.1101/566000

**Authors:** Amanda Thomaz, Kelly de Vargas Pinheiro, Bárbara Kunzler Souza, Lauro Gregianin, Algemir L. Brunetto, André T. Brunetto, Caroline Brunetto de Farias, Mariane da Cunha Jaeger, Vijay Ramaswamy, Carolina Nör, Michael D. Taylor, Rafael Roesler

**Author notes:** Correspondence: Rafael Roesler, Department of Pharmacology, Institute for Basic Health Sciences, Federal University of Rio Grande do Sul, Rua Sarmento Leite, 500 (ICBS, Campus Centro/UFRGS), 90050-170 Porto Alegre, RS, Brazil.

## Abstract

Neurotrophins are critically involved in regulating normal neural development and plasticity. Brain-derived neurotrophic factor (BDNF), a neurotrophin that acts by binding to the tropomyosin receptor kinase B (TrkB) receptor, has also been implicated in the progression of several types of cancer. However, its role in medulloblastoma (MB), the most common type of malignant brain tumor afflicting children, remains unclear. Here we show that selective TrkB inhibition with the small molecule compound ANA-12 impaired proliferation and viability of human UW228 and D283 MB cells, and slowed the growth of MB tumors xenografted into nude mice. These effects were accompanied by increased apoptosis, reduced extracellular-regulated kinase (ERK) activity, increased expression of signal transducer and activator of transcription 3 (STAT3), and differential modulation of p21 expression dependent on the cell line. In addition, MB cells treated with ANA-12 showed morphological alterations consistent with differentiation, increased levels of the neural differentiation marker β-III Tubulin (TUBB3), and reduced expression of the stemness marker Nestin. These findings are consistent with the possibility that selective TrkB inhibition can display consistent anticancer effects in MB, possibly by modulating intracellular signaling and gene expression related to tumor progression, apoptosis, and differentiation.

## INTRODUCTION

Medulloblastomas (MBs) are highly aggressive and heterogeneous brain tumors of the cerebellum that account for about 20% of pediatric brain cancers (Ramaswamy and Taylor, 2017; Northcott et al., 2019). Integrative genomic and epigenomic studies on MB biology classify this disease into four clinically relevant consensus subgroups: wingless (Wnt), Sonic hedgehog (Shh), group 3 and group 4 (Northcott et al., 2012; Taylor et al., 2012). The 2016 World Health Organization (WHO) Classification of Tumors of the Central Nervous System recently acknowledged the molecular subgroups in the classification of MBs, providing clinical utility for the improvement of MB diagnosis (Louis et al., 2016). More recent advancements in deep transcriptional and methylation profiling of 763 primary MB samples revealed new subtypes within each of the four subgroups and further classified MB into 12 subtypes: two Wnt, four Shh, three group 3, and three group 4 subgroups (Cavalli et al., 2017). Current treatment is based in risk stratification and despite significant advances in surgery, radiotherapy, and drug therapy, effective treatment to MB remains a challenge (Kumar et al., 2017; Northcott et al., 2019). Because irradiating the central nervous system (CNS) can be harmful to the developing brain, radiation therapy is typically avoided in children under the age of three, but this can compromise disease control and survival. Consequently, there is a need for new treatments that can be tolerated in the younger population to treat therapy-resistant disease as well as to decrease potential side effects (Sabel et al., 2018; Northcott et al., 2019).

Targeting oncogenic fusions and dysregulated signal transduction pathways is an approach that can potentially improve the outcome of pediatric tumors therapy. Neurotrophins (NT) are a group of growth factors that stimulates cell survival pathways through the activation of the tropomyosin receptor kinase (Trk) receptors. The Trk family includes TrkA, TrkB and TrkC, which are encoded by NTRK1, NTRK2 and NTRK3 genes, respectively. TrkA is the high-affinity receptor for nerve growth factor (NGF), whereas TrkB has high affinity for brain-derived neurotrophic factor (BDNF) and NT-4. NT-3 can bind to all Trk receptors but has highest affinity for TrkC and is the sole ligand of this receptor (Chao, 2003; Park and Poo, 2013). NTs were initially characterized for their roles in regulating development and plasticity in the CNS (Park and Poo, 2013). However, it is now clear that deregulation of NT signaling is involved in the pathogenesis and progression of several tumor types (Vaishnavi et al., 2015). Gene fusions involving NTRK genes that lead to transcription of chimeric Trk proteins with constitutively activate or overexpressed kinase function conferring oncogenic potential have become increasingly important targets for cancer therapy (Cocco et al., 2018; Lange and Lo, 2018).

Expression of NTs and Trks has been reported in MB tumor samples from patients (Washiyama et al., 1996), and activation of different Trk receptors can influence MB cell function (Chou et al., 1997). Stimulation of TrkA by NGF and TrkC by NT-3 typically increases cell death and differentiation (Kim et al., 1999; Li et al., 2010; 2016), and higher TrkC expression has been associated with a favourable outcome and predictor of survival of MB patients (Segal et al., 1994; Friedrich et al., 2017). Despite substantial knowledge about TrkA and TrkC receptors in MB, the role of BDNF/TrkB in MB cells remains poorly understood (Venkatesh H and Monje, 2017). Previous reports showed that BDNF or TrkB inhibition can display either pro- or antitumoral effects in these tumors (Schmidt et al., 2010; Nör et al., 2011; Thomaz et al., 2016). Given that BDNF/TrkB pathway is implicated in the pathogenesis and prognosis of a wide variety of malignancies, including neuroblastoma, glioblastoma, head and neck, breast, lung, and pancreas tumors as well as leukemia, being associated with promotion of proliferation, migration, resistance to anoikis and resistance to chemotherapy (Thiele et al., 2009; Roesler et al., 2011; Radin and Patel, 2017; Meng et al., 2019), here we investigated the potential role of TrkB inhibition in experimental MB.

## MATERIALS AND METHODS

### Ethics Statement

All experimental procedures were performed in accordance with the Brazilian Guidelines for the Care and Use of Animals in Research and Teaching (DBCA, published by CONCEA, MCTI; https://www.mctic.gov.br/mctic/opencms/institucional/concea/paginas/concea.html), and approved by the institutional Animal Care Committee (*Comissão de Ética no Uso de Animais-CEUA, Hospital de Clínicas de Porto Alegre-HCPA*), under protocol number 20160098.

### Cell Culture

Human MB cell lines D283 and UW-228 were originally obtained from the American Type Culture Collection (ATCC, Rockville, USA). These two cell lines present molecular features of different MB molecular subgroups: UW228 cells are TP53-mutated and classified as Shh, whereas D283 cells are p53 wild-type and classified as Group ¾ (Ivanov et al., 2016). The D283 cell line was cultured in Dulbecco’s modified Eagle’s medium (DMEM low glucose, Gibco, Grand Island, USA) while UW228 cell line was cultured in DMEM: Nutrient Mixture F-12 (DMEM/F-12 Gibco^®^), both medium supplemented with 10% (v/v) fetal bovine serum (FBS, Gibco) and 1% (v/v) penicillin/streptomycin (Gibco). Cells were incubated in a humidified atmosphere of 5 % CO^2^ at 37 °C.

### Drug Treatments

ANA-12 (Sigma Aldrich, St. Louis, USA) was dissolved in dimethyl sulfoxide (DMSO, Sigma Aldrich, MO, USA) at the concentration of 6140 μM. BDNF (Sigma Aldrich, USA) was diluted in sterile ultrapure water at the stock solution of 1000 ng/ml. Cells were treated with increasing concentrations of ANA-12 (5, 10, 20, or 30 μM) or human recombinant BDNF (50 ng/ml) in complete medium for 6, 24, 48 or 72 h. The concentration of the vehicle DMSO was used as control and did not exceed 0.5 % (v/v).

### Mice and *In Vivo* Experiments

*In vivo* studies were performed in accordance with procedures approved by the Brazilian Guidelines for the Care and Use of Animals in Research and Teaching (DBCA, published by CONCEA, MCTI) and approved by the institutional Animal Care Committee (CEUA-HCPA) under protocol number 160098.

Balb/c nude mice males and females (6 to 12 weeks old) were kept under aseptic conditions in ventilated cages and received food and water ad libitum. A total of 1 × 10^6^ D283 cells were processed in serum-free DMEM and diluted 1:1 with Matrigel (Corning, Corning, USA). Two-hundred microliters of cell suspension was subcutaneously inoculated into the lower right dorsum of nude mice generating xenograft tumors. When established tumors reached approximately 80 to 100 mm^3^, the mice were randomly divided into two groups (12 mice/group) and subjected to treatments. The TrkB selective inhibitor ANA-12 was dissolved in DMSO at 6.5 mg/ml and administered at 1 mg/kg (ANA-12 + saline solution + Tween 20 2%) or vehicle alone (DMSO 2% + saline solution + Tween 20 2%) once daily by intraperitoneal (IP) injections for 15 days. The drug dose dose was chosen on the basis of previous studies using systemic administration of ANA-12 (Cazorla et al., 2011; Chodroff et al. 2016). After treatment, all the mice were euthanized, and the tumors were excised, measured and weighed. The tumor size for the xenografts was determined using a calliper, and the volume was calculated: tumor volume (mm3) = [(Width 2 × Length)/2], where the width is the smallest measurement and the length is the longest measurement.

### Cell Viability

MB cells were seeded at density of 3×10^3^ cells per well in complete medium into 96-well plates (NEST Biotechnology, Jiangsu, China). After overnight culture in complete medium, cells were treated with ANA-12 as described above and after 24, 48 and 72 h of treatment, the medium was removed, cells were washed with phosphate-buffered saline (PBS), and 50 μl of 0.25 % trypsin/EDTA (Gibco by Life Technologies) solution was added to detach cells. To assess cell viability, the cell suspension was homogenized with 0.4 % trypan blue 1:1 and counted immediately in a hemocytometer. Experiments were performed at least four times in quadruplicates for each treatment. Cell viability was normalized to the control DMSO. For the calculation of IC_50_, data were fitted in a dose response curve (Graphpad Prism v. 6.0) using the equation: y=min+(max − min)/(1 + 10^Λ^((LogIC_50_ − x)*Hillslope + Log ((max −min)/(50-min) − 1))).

### Ki67 Proliferation Assay

The Muse Ki67 Proliferation kit (Merck, Princeton, USA) was used to detect proliferating and non-proliferating cells based on Ki67 expression. MB cells were plated at 2 × 10^5^ cells per well in 6-well plate (NEST) and treated with ANA-12 for 24 h. After treatment, the supernatant was removed, cells were detached, counted and adjusted to the concentration of 1 × 10^5^ cells, followed by washing, fixation, permeabilization and centrifugation steps. Cells were stained with anti-Ki67-PE antibody or IgG1-PE antibody (isotype), used as negative control, for 30 min, in the dark, at room temperature according to the manufacturer’s instructions. Percentage of Ki67 negative and positive cells was determined from the fluorescence of cells in each sample analysed by Muse Cell Analyzer (Merck). Experiments were performed at least four times in duplicates for each treatment.

### Apoptosis Assay

The Annexin V-FITC apoptosis detection kit (BD Biosciences, San Diego, USA) was used to detect apoptosis and cell death, respectively. MB cells were plated at 15 × 10^3^ cells per well in 24-well plate (NEST) and cells were treated with ANA-12 or BDNF for 24 and 48 h. After treatment times, both floating and attached cells were harvested and washed twice with ice-cold PBS, resuspended in 1× binding buffer, and stained with Annexin V-FITC and PI for 15 min, in the dark, at room temperature. Percentage of Annexin V-FITC-positive and PI-positive cells was determined from the fluorescence of 20.000 events for each sample in a flow cytometer (Attune Acoustic focusing cytometer, Applied Biosystems, Beverly, USA). Data was analyzed using Attune Cytometric Software version 1.2.5. At least 3 independent replicates were performed.

### PI3K and MAPK Dual Pathway Activation Assay

MB cells were plated at 2 × 10^5^ cells per well in 6-well plate (NEST) and treated with ANA-12 for 24 h. To access the activation of both PI3K and MAPK signaling pathways the Muse^™^ PI3K/MAPK Dual Pathway Activation Kit (Merck) was used. After treatment, the supernatant was removed, cells were detached, counted and adjusted to the concentration of 1 × 10^5^ cells, followed by washing, fixation, permeabilization and centrifugation steps. Cells were stained with anti-phospho-Akt (Ser473) conjugated with Alexa Fluor-555 and anti-phospho-ERK1/2 (Thr202/Tyr204 and Thr185/Tyr187) conjugated with PECy5 for 30 min, in the dark, at room temperature according to the manufacturer’s instructions. Percentage of phospho-AKT and phospho-ERK positive cells were determined from the fluorescence of cells in each sample analysed by Muse Cell Analyzer (Merck). Experiments were performed at least four times in duplicates for each treatment.

### mRNA Expression

Analysis of mRNA expression was performed in MB cell lines seeded at density of 1.8 × 10^6^ cells in T75 cm^2^ culture flasks (NEST) and treated with ANA-12 or control vehicle for 6 or 24 h. After treatment period, cells were counted and adjusted to the concentration of 1 × 10^6^. Total RNA purification was performed using the kit SV total RNA isolation system (Promega, Fitchburg, USA). Purified total RNA was quantified using NanoDrop (Thermo Fisher Scientific, Wilmington, USA) and 500 ng of total RNA was used to generate cDNA using GoScript Reverse Transcriptase kit (Promega), according to the manufacturer’s instructions. mRNA expression levels of target genes (p21, STAT3, Nestin and TUBB3) were performed using real-time reverse transcriptase (RT-PCR) with SYBR Green master mix (Applied Biosystems) and analyzed by StepOnePlus Real-Time PCR System (Thermo Fisher Scientific). Cycling was performed as follows: 50°C for 2 minutes and 95°C for 10 minutes, followed by 40 cycles of 95°C for 15 seconds and 60°C for 30 seconds. This was followed with a dissociation stage of 95°C for 15 seconds, 60°C for 30 seconds, and 95°C for 15 seconds. Samples were analysed and calculated using the ΔCT method from triplicate reactions, with the levels of gene normalized to the relative Ct value of GAPDH. mRNA expression levels of target genes (BDNF and TrkB) were performed using conventional PCR. Experiments using PCR were performed using GoTaq Hot Start Polymerase (Promega) according with manufacturer’s instruction. Cycling conditions were as follows: 35 cycles for amplification that consisted of 1 min at 95 °C, denaturation at 94 °C for 30 s, annealing at 60 °C, for 30 s, and extension of primers at 72 °C for 45 s, followed by a final extension at 72 °C for 10 min. The products of BDNF, TrkB, and GAPDH were electrophoresed through 1.5 % agarose gels containing ethidium bromide (Biotium, Fremont, CA, USA) and visualized with ultraviolet light. Primers are listed in Table 1.

**TABLE 1.**
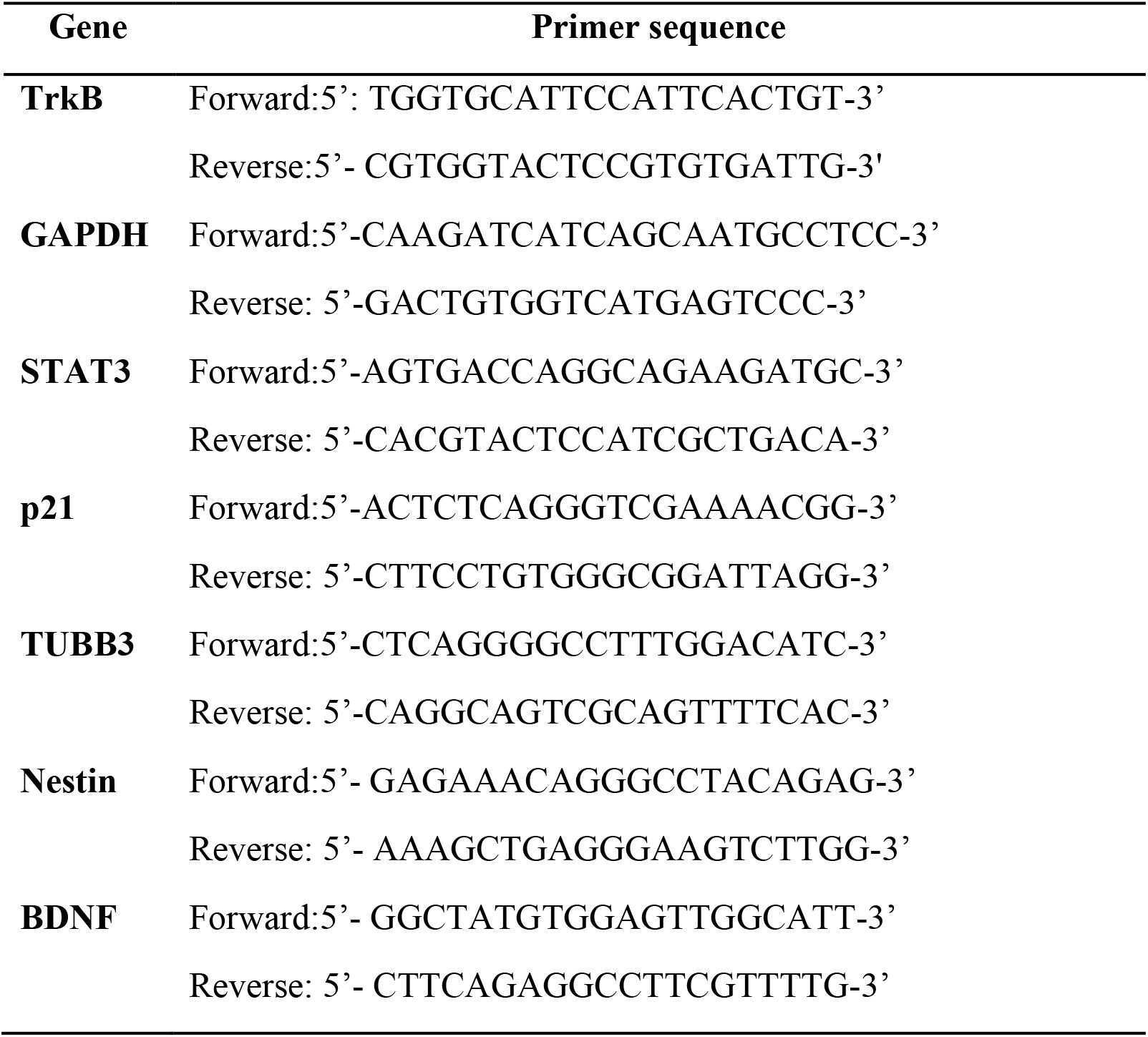
Forward and reverse primer sequences used for RT-PCR amplification

### Protein Extraction and Immunoblotting

MB cells were plated at density of 1.8 × 10^6^ cells in T75 cm^2^ culture flasks (NEST) and treated with ANA-12 or control vehicle for 6 or 24 h. After treatment, cells were lysed in cell lysis buffer (Cell Signaling Technology, Danvers, USA) containing protease and phosphatase inhibitors (Sigma Aldrich) and centrifuged at 10,000 rpm for 10 minutes at 4°C. Protein concentration was determined by Bradford reagent (Bio-Rad, Hercules, USA) using the spectrophotometer SpectraMax Plus 384 Microplate Reader (Molecular Devices, San Jose, USA) at wavelength of 540 nm. Subsequently 20 μg of protein were separated by SDS-polyacrylamide gel electrophoresis and transferred onto a polyvinylidene difluoride membranes (Immobilon-P PVDF, EMD, Merck,). Membranes were blocked with 5% fat-free milk in TBST (0.01% Tween) for 1 h, incubated with antibodies against p21(1:200, Santa Cruz Biotechnology, Santa Cruz, USA) and β-Tubulin III (1:1000, Abcam, Cambridge, MA, USA) overnight at 4°C and followed by 1-h incubation at room temperature with HRP-conjugated anti-rabbit secondary antibody (1:2000, Sigma Aldrich). Chemiluminescence was detected using ECL Western Blotting substrate (Merck, USA) and analysed by ImageQuant LAS500 (GE Healthcare Life Sciences, Little Chalfont, UK). Membrane was stained with Coomassie blue (0.025%) for protein load control. Densitometric analyses were performed using ImageJ software and relative densitometric unit (RDU) was calculated by the normalization of interest protein level to Coomassie blue staining. ANA-12 treated cells were corrected by control groups (DMSO treated cells). Three individual replicates were performed.

### Immunohistochemistry

ANA-12 and vehicle-treated D283 xenografts samples were harvested, formalin-fixed, paraffin-embedded and sectioned at four microns. Antigen retrieval was performed using preheated pH 6.0 citrate buffer. Endogenous peroxide activity was blocked using 5% H2O2 for twenty minutes. Sections were blocked and stained with primary antibodies against: Ki67 (1:200, Abcam), p53 (1:100, Santa Cruz), TrkB (1:800, Abcam) and phospho-TrkB (Y706/707, 1:100, Abcam) overnight at 4°C. Secondary antibody incubation was performed at room temperature for 90 min utilizing a horseradish-peroxidase (HRP)-labeled anti-rabbit IgG (Merck), and staining was visualized with DAB (Dako, CA, USA) following manufacturer’s instructions. All slides were counterstained with hematoxylin, dehydrated and permanently mounted using standard procedures. Quantitative evaluation was made using IHC profiler plugin in the software image J (Varghese et al., 2014).

### Statistical Analysis

All statistical analyses were performed using GraphPad Prism software version 6.0. Data were represented as means ± standard error of mean (SEM). For comparison between two data sets, a two-tailed unpaired Mann-Whitney test was used. For analysis of three or more sets of data, ANOVA test followed by Bonferroni multiple comparison test was used; *p* < 0.05 was considered to indicate statistical significance.

## RESULTS

### Selective TrkB Inhibition Suppresses Cell Viability and Proliferation of MB Cells

To evaluate the effects of TrkB inhibition on MB cell viability, we exposed the cells to varying concentration of ANA-12 (5, 10, 20 or 30 μM) for different time periods (24, 48 or 72 h). A dose-dependent reduction of cell viability was observed in both cell lines, and increasing effects were observed with longer exposure ANA-12 (**Figure 1A**). Fifty percent inhibition of growth (IC_50_) was calculated for ANA-12 considering different exposure times, ranging from 23.99 to 17.42 μM in UW228 cells and 24.83 to 14.74 in D283 cells (**Figure 1B**). Moreover, changes in cell viability were accompanied by alterations in cell morphology characteristic of cell death features in both cell lines, and differentiation in UW228 cells (**Figure 1C**).

**FIGURE 1.**
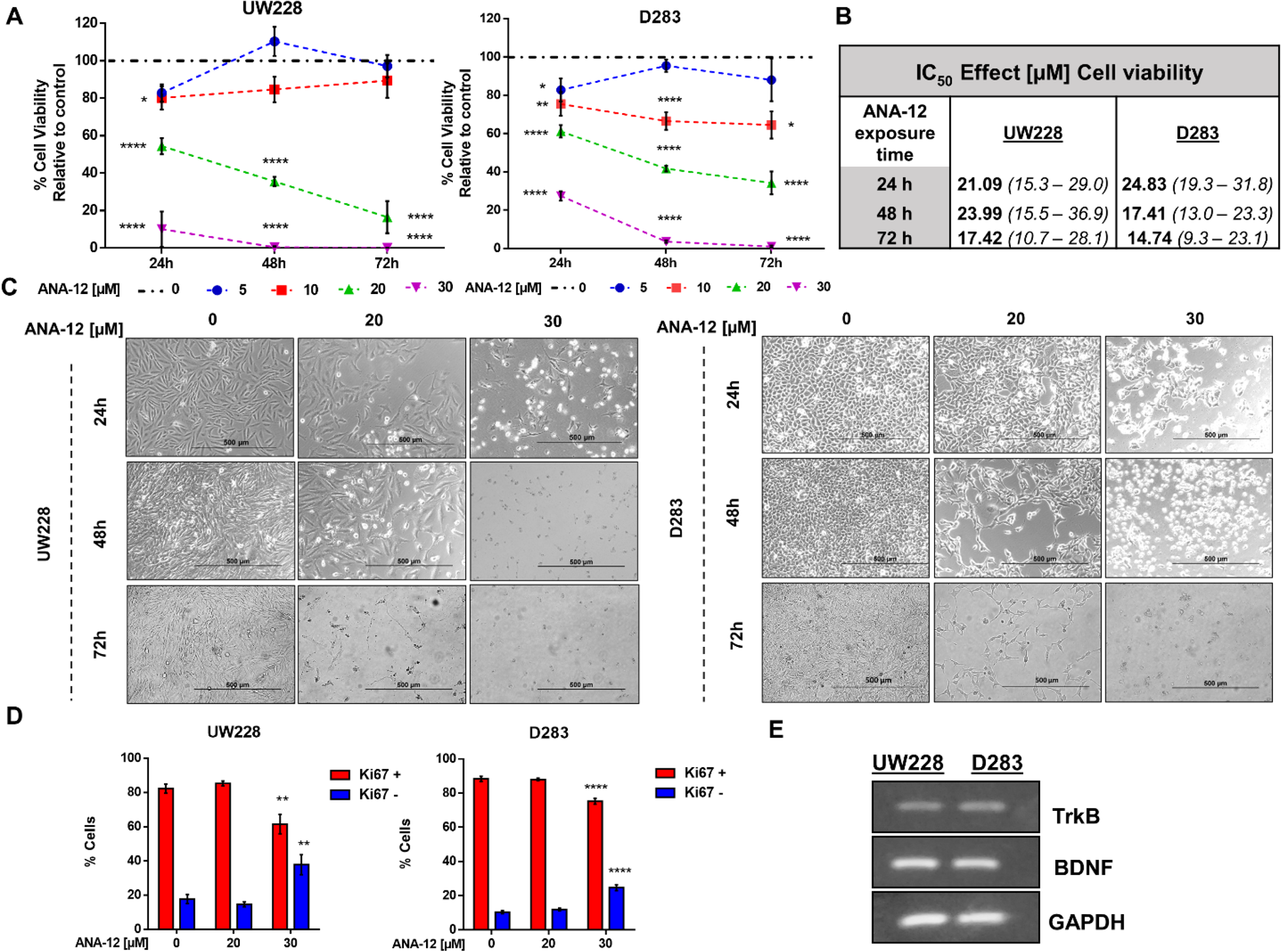
TrkB inhibition reduces MB cell viability and proliferation in a dose- and time-dependent manner. D283 and UW228 MB cells lines were treated with increasing concentrations of ANA-12. The vehicle DMSO served as control. (**A**) After 24, 48 or 72 h of ANA-12 exposure, cell viability was assessed by trypan blue cell counting; *n* = 4 independent experiments. (**B**) Comparison of the half inhibitory concentration (IC50) for ANA-12 in MB cells. IC50 values are represented as the IC50 best fit value (μM) and therange of 90% interval of confidence. (**C**) Representative morphology of cell cultures under different treatment concentrations at 24, 48 and 72 h. The left panel displays UW228 cells and right panel D283 cells. The reduction in cell density and change in cell morphology is showed. Scale bar 500 μm. (**D**) After 24 h of ANA-12 exposure, proliferation of UW228 (left panel) and D283 (right panel) cells was measured by staining with the Ki67 marker and analysed by Muse Cell Analyzer; *n* = 4 independent experiments. (E) RT-PCR confirming mRNA expression of TrkB, BDNF and GAPDH in MB cells. Data are shown as mean ± SEM; \**p* < 0.05, ** *p* < 0.01, *** *p* < 0.001, \*\*\*\**p* < 0.0001 compared to controls.

We next assessed cell proliferation using the Ki67 marker in cells exposed to ANA-12 for 24 h. Exposure to ANA-12 for 24 h resulted in a significant decrease in the percentage of cells positive for Ki67 (**Fig. 1D**). Expression of BDNF and TrkB in MB cells was confirmed by PCR analysis (**Figure 1E**). These results indicate that TrkB inhibition reduces viability and proliferation in cell lines representative of different molecular subgroups of MB.

### TrkB Inhibition Impairs MB Growth in Mice D283 Xenograft with D283 MB Tumors

We next assessed the effects of ANA-12 on MB growth *in vivo*. D283 MB cells were grown subcutaneously in nude mice until tumors were apparent (~80-100 mm^3^). ANA-12 was administered i.p. at 1mg/kg daily for 15 days. Control animals received vehicle alone (DMSO 2%) in the same injection regimen. The tumor growth was significantly delayed in ANA-12 treated mice compared with vehicle-treated controls (**Figure 2**). Mice treated with ANA-12 showed a slower rate of tumor growth at 12 and 15 days of treatment and an average reduction in tumor volume of 35%, whereas growth continued at a high rate in control animals (**Figure 2B**). *Ex-vivo* tumor analysis showed an apparent reduction in tumor volume and weight, however these differences did not reach statistical significance (**Figure 2C, 2D, 2E**). Importantly, the treatment protocol used did not induce overt toxicity and there were no significant changes in animal weights over the study period (**Figure 2H**). We analysed the protein content of TrkB and phospho-TrkB in tumor samples and found reduced levels of phospho-TrkB in tumors from mice treated with ANA-12. The antibody used detects the double specific residues Y706/707 that are positioned at the autophosphorylation site, which is essential for the TrkB activation. The phosphorylation of these specific sites also recruits adaptor proteins when phosphorylated, including Grb2 and SH2B, that can participate in the signalling through the ERK and AKT pathways (Boltaev et al. 2017; Saarelainen et al. 2003). Ki67 IHC staining was used to investigate the extent of proliferation in the tumors. Tumors from mice receiving ANA-12 showed a reduction of approximately 20% in Ki67-positive tumor cells. Finally, we verified whether ANA-12 would affect p53 expression. There was no difference in p53 staining between ANA-12-treated and control groups (**Figure 2F, 2G**). These results indicate that TrkB inhibition results in significant *in vivo* antitumoral activity in MB xenografts.

**FIGURE 2.**
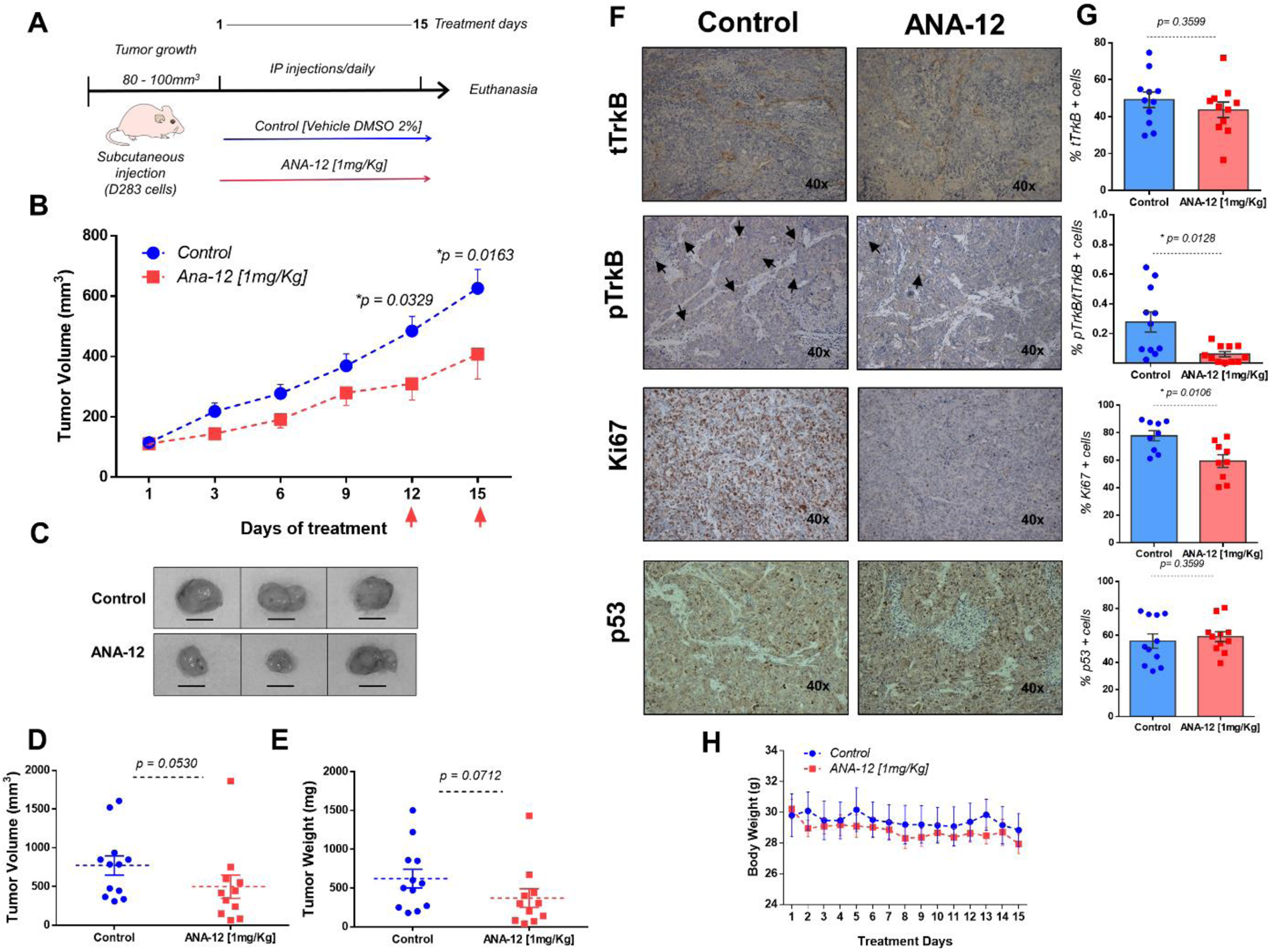
ANA-12 delays tumor growth in nude mice xenografted with MB cells. (**A**) A total of 1 × 10^6^ D283 cells were injected subcutaneously into the right flank and the mice were analysed for tumor growth by manual calliper rule. After tumors reached the volume of 80-100 mm^3^, the mice were treated intraperitoneally with 1 mg/kg of ANA-12 once daily for 15 days. Control animals received DMSO 2%. (**B**) Tumors were measured every two days and volumes were calculated as described in Materials and Methods. Tumor growth is represented by tumor volume (mm^3^) at the indicated days; ANA-12, *n* = 11, and DMSO, *n* = 12. (**C**) Representative macroscopic appearance of representative tumors from mice injected with DMSO (upper panel) or ANA-12 (lower panel). (**D**) Tumor volumes (mm^3^) at the time of tissue harvest. (**E**) Tumor weight (mg) at the time of tissue harvest. (**F**) Representative images of TrkB total (tTrkB), phospho-TrkB (pTrkB), Ki67 and p53 staining of MB tumors from mice given DMSO and ANA-12. (**G**) Percentage of tTrkB, pTrkB, Ki67 and p53 positivity in tumors from mice treated with either DMSO 2% or ANA-12. (**H**) Mice were weighed daily; mean ± SEM body weights (g) across treatment days. Data are expressed by mean ± SEM. Statistically significant differences found between ANA-12 treatment and control are indicated by the *p* values shown in the figure.

### TrkB Inhibition Induces Pro-Apoptotic Effects in MB Cells

We assessed the induction of apoptosis in cells treated with different doses of ANA-12 or recombinant BDNF (50 ng/ml) for 24 and 48 h by flow cytometry analysis of Annexin V/PI staining. A significant increase in cell death was observed in response to ANA-12 at 30 μM. Also, there was an increase in the percentage of cells positive for Annexin V after 24 h of treatment, when compared with either control or BDNF-treated cells, 38.67% of UW228 cells (**Figure 3A, 3C**) and 28.78% for D283 cells (**Figure 3B, 3D**) were stained for Annexin V only and were considered apoptotic cells. After 48 h of treatment, there was an increase in the percentage of cells stained for both Annexin V and PI (dead cells), namely 44.84% in UW228 cells and 17.36% in D283 cells. Thus, TrkB inhibition dose-dependently induced apoptosis.

**FIGURE 3.**
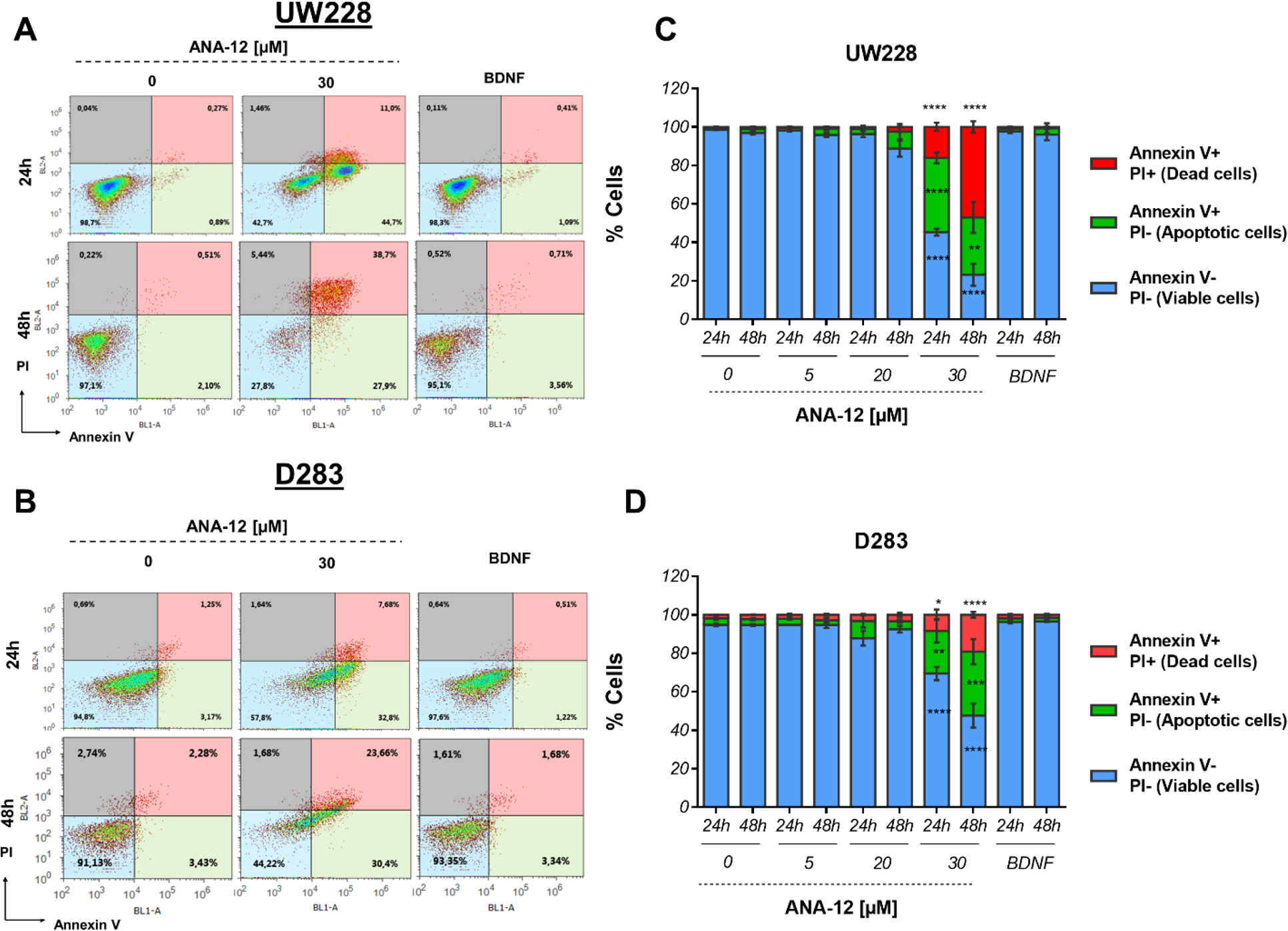
Proapoptotic effects of TrkB inhibition in MB cells. UW228 and D283 cells were stained with Annexin V and propidium iodide (PI) after treatment with ANA-12 (5, 20 and 30 μM) or BDNF (50ng/ml) for 24 or 48 h. The vehicle DMSO served as control. (**A, B**) Representative density plots of flow cytometry of UW228 (upper panel) and D283 (lower panel) cells. (**C, D**) Percentages of Annexin V/PI apoptosis assays. Viable cells are negative for Annexin V and PI, apoptotic cells are positive for Annexin V and negative for PI, and dead cells are positive for both Annexin V and PI; *n* = 3 independent experiments; UW228 (upper panel) and D283 (lower panel). Data are expressed as mean ± SEM; \**p* < 0.05, \*\**p* < 0.01, \*\*\**p* < 0.001, and \*\*\*\**p* < 0.0001 compared to controls.

### The Antiproliferative and Proapoptotic Effects of TrkB Inhibition are Associated with Extracellular-Regulated Kinase (ERK) Pathway Regulation

To investigate molecular pathways associated with the antiproliferative and proapoptotic effects of TrkB inhibition, we evaluated the activation of two major pathways simultaneously, protein kinase B (PKB or AKT) and extracellular-regulated kinase (ERK), which are involved in mediating intracellular responses to TrkB activation. After exposure of 30 μM ANA-12 for 24 h, a decrease in the percentage of phospho-ERK positive cells was detected in UW228 cells (15.46%) in comparison with control cells (32.25%) (**Figure 4A, 4B**, left panels). In D283 cells, ANA-12 at 20 μM or 30 μM was effective in reducing phospho-ERK (6.16% and 4.16%, respectively) when compared with control cells (16.94%) (**Figure 4A, 4B**, right panels). There was also increases in negative cells, and no differences were observed for AKT or dual-pathway activation.

**FIGURE 4.**
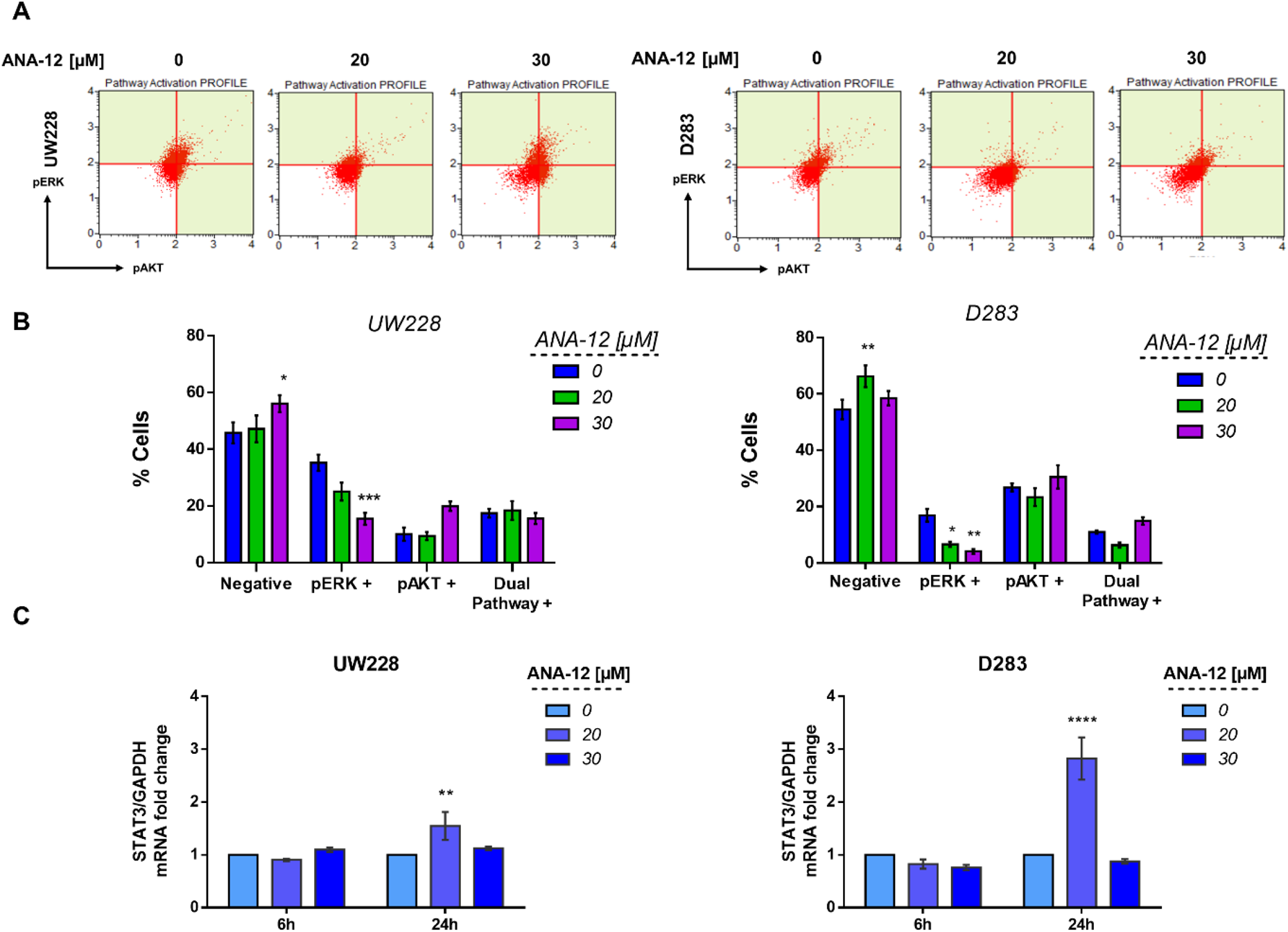
TrkB inhibition decreases ERK activation and increases the expression of STAT3 mRNA levels in MB cells. (**A**) Representative dot plots of UW228 (right) and D283 (left) cells treated with ANA-12 for 24 h. MB cells were stained with pAKT or pERK antibodies and analysed by Muse Cell Analyzer. (**B**) Percentage of cells are represented as follows: negative cells are negative for pERK and pAKT; pERK+ cells are positive for pERK marker; pAKT+ cells are positive for pAKT marker and dual pathway + cells are positive for pERK and pAKT markers; *n* = 3 independent experiments. (**C**) STAT3 mRNA levels in MB cells after ANA-12 exposure for 6 or 24 h. Gene expression levels were normalized by GAPDH level and corrected by control; *n* = 3 independent experiments. Data are shown as mean ± SEM; * *p* < 0.05, ** *p* < 0.01, *** *p* < 0.001, \*\*\*\**p* < 0.0001 compared to controls.

We also evaluated the transcriptional expression level of the Signal transducer and activator of transcription 3 (STAT3), which is a known downstream mediator of BDNF/TrkB, AKT and ERK signaling. MB cells exposed to ANA-12 at 20 μM for 24 h, concentrations close to IC_50_ values, resulted in approximately 1.5 and 2.8-fold change increases of STAT3 transcriptional expression in UW228 and D283 cells respectively (**Figure 4C**). These results support the possibility that reductions in ERK activation may be involved in the antiproliferative and proapoptotic actions of TrkB inhibition in MB cells, and the treatment also results in increased expression of STAT3.

### TrkB Inhibition Regulates p21 Gene Expression in MB Cells

The cyclin-dependent kinase (CDK) inhibitor p21 is a well-known tumor suppressor that promotes cell cycle inhibition, transcriptional regulation, and modulation of apoptosis. In order to access the status of p21 expression after TrkB inhibition, we treated MB cells with ANA-12 for 6 or 24 h and then performed RT-qPCR and Western blot analyses. TrkB inhibition led to increases in both mRNA expression and protein levels of p21 in UW228 cells in a dose-dependent manner. In addition, we observed 3 a 10.8-fold change increase of p21 mRNA expression in UW228 cells after 6 and 24 h of ANA-12 respectively (**Figure 5A**, left panel). UW228 cells exposed to 20 and 30 μM of ANA-12 for 6 h showed approximately 1.97 and 2.49-fold change increases in the protein levels of p21 (**Figure 5B, 5C**, left panel).

**FIGURE 5.**
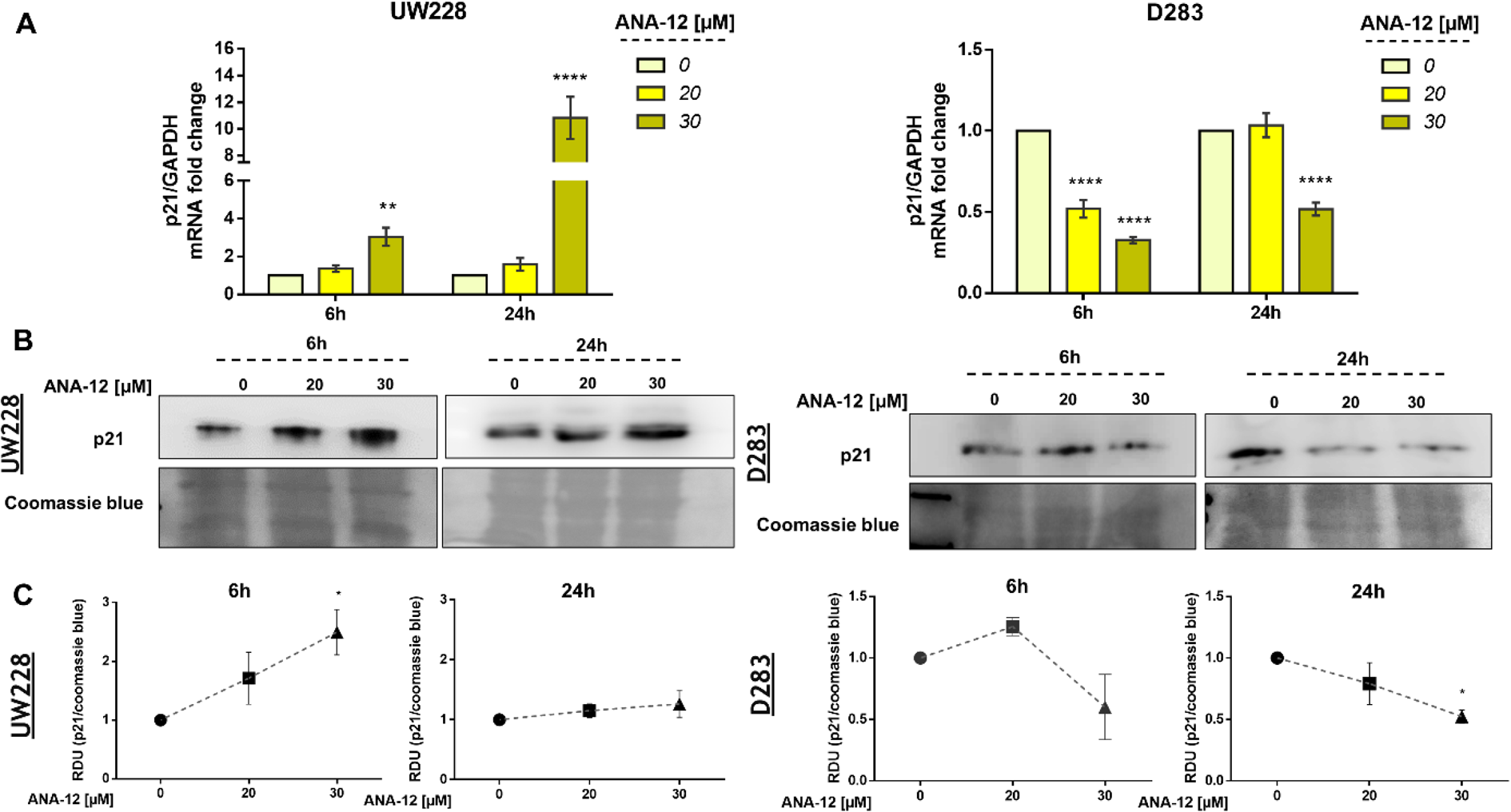
TrkB inhibition promotes changes p21 gene expression in MB cells. (**A**) p21 mRNA levels in UW228 (left panel) and D283 (right panel) cells after ANA-12 exposure for 6 or 24 h. Gene expression levels were normalized by GAPDH level and corrected by control; *n* = 3 independent experiments. (**B**) Western blot of p21 protein levels in UW228 (left panel) and D283 (right panel) cells after ANA-12 exposure for 6 or 24 h; *n* = 3 independent experiments. (**C**) Relative densitometric unit (RDU) analyses of p21 protein levels were normalized by Coomassie blue (loading control) and corrected by control; UW228 (left panel) and D283 (right panel); *n* = 3 independent experiments. Data are shown as mean ± SEM; \**p* < 0.05, \*\**p* < 0.01, \*\*\**p* < 0.001, \*\*\*\**p* < 0.0001 compared to controls.

Interestingly, opposite effects of TrkB inhibition on p21 expression were found in in D283 cells. Significant 2 to 3.3-fold decreases in p21 mRNA levels were observed in D283 cells treated with ANA-12 for 6 and 24 h respectively (**Figure 5A**, right panel). We also detected a 2.5-fold change decrease in p21 protein content in D283 cells exposed to ANA-12 for 24 h (**Figure 5B, 5C**, right panel). These results indicate differential responses of p21 expression to TrkB inhibition in different cell lines.

### Inhibiting TrkB Induces Morphological Changes and Alters the Expression of Differentiation and Pluripotency Markers

We went on to investigate whether morphological changes observed after ANA-12 treatment would be associated with changes in differentiation in MB cells. We observed neurite-like extensions after 24 h of ANA-12 treatment, particularly in UW228 cells (**Figure 6A**, left panel) whereas D283 cells showed predominantly morphologic features associated with cell death (**Figure 6A**, right panel). Next, we performed flow cytometry analysis to evaluate the forward and side scatter characteristics, particularly the SSC-A parameter that indicates cell granularity or internal complexity. We observed an increase in cell complexity and granularity in UW228 cells treated with ANA-12 at 30 μM for 24 h (**Figure 6B, 6C**). Moreover, we measured the expression level of the neural differentiation marker β-III Tubulin (TUBB3) and the neural stem/progenitor cell marker Nestin. We found that the transcriptional expression level of Nestin was decreased in dose-dependent manner in both MB cell lines (**Figure 6D**). We also observed an 1.8-fold change increase in TUBB3 mRNA levels in UW228 cells (**Figure 6E**), however, there were no differences in the protein content of TUBB3 (**Figure 6F, 6G**). These findings suggest that UW228 cells, classified SHH MB subtype, display features of cell differentiation after TrkB inhibition, and both cell lines showed reductions of the pluripotency marker Nestin.

**FIGURE 6.**
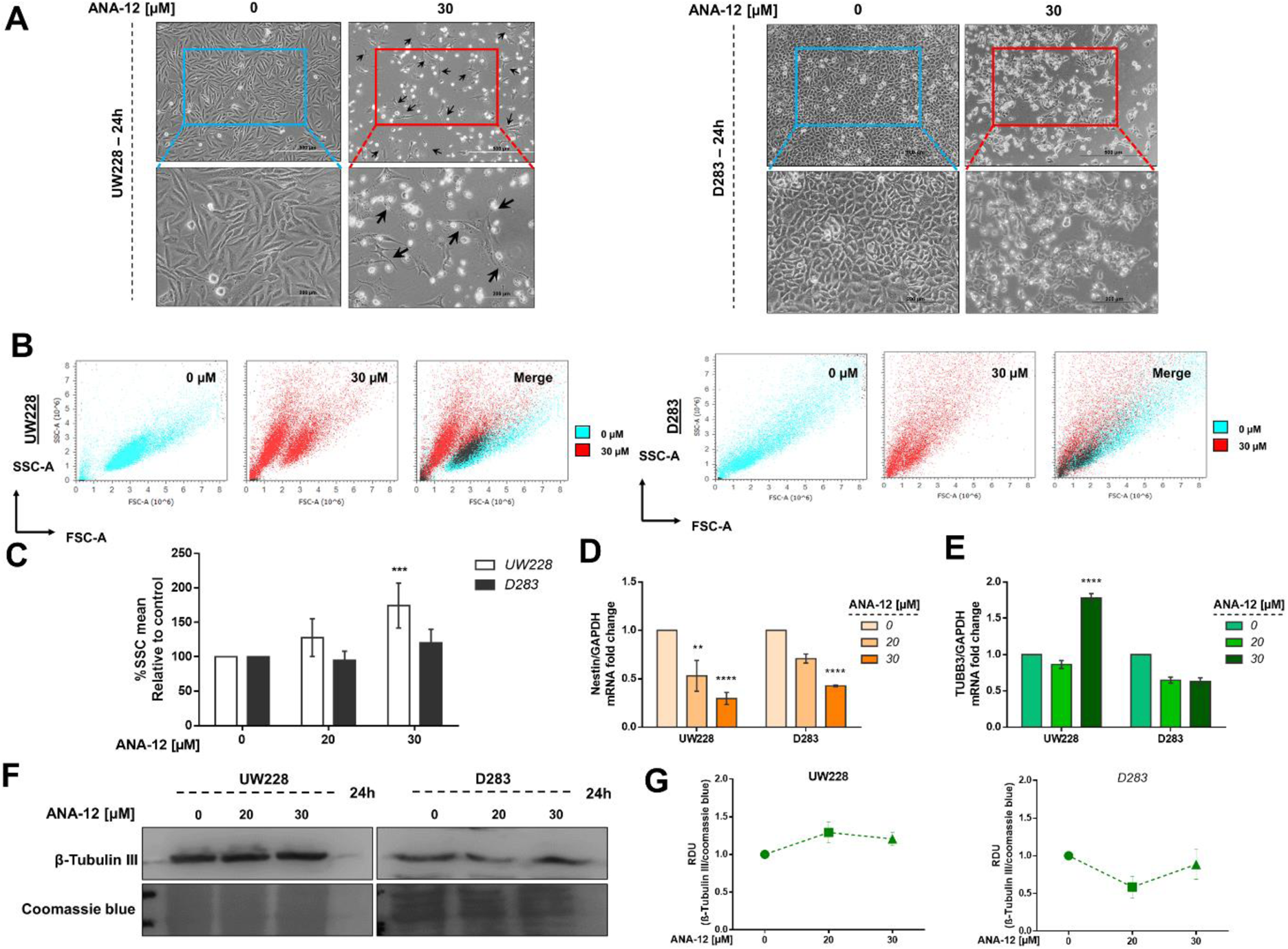
TrkB inhibition increases cell complexity and differentiation in and decreases pluripotency. (**A**) Morphology of UW228 (left panel) and D283 (right panel) cells after ANA-12 exposure for 24 h. Black arrows represent neurite-like extensions. Upper panel scale bar: 500 μM and lower panel scale bar: 200 μM. (**B**) Representative dot plots of UW228 (lower) and D283 (right) cells treated with ANA-12 for 24 h and analysed by flow cytometry. Cyan dots show control cells, red dots show ANA-12 treated cells and black dots in merge graphics show overlay of control and treated cell populations. (**C**) Percentages of SSC-A mean from MB cells exposed with the indicated concentrations of ANA-12 for 24 h; *n* = 4 independent experiments. (**D**) Nestin mRNA levels in MB cells after ANA-12 exposure for 24 h. (**E**) TUBB3 mRNA levels in MB cells after ANA-12 exposure for 24 h. Gene expression levels were normalized by GAPDH level and corrected by control; *n* = 3 independent experiments. (**F**) Western blot of β-Tubulin III protein levels in MB cells after ANA-12 exposure for 24 h; *n* = 3 independent experiments. (**G**) Relative densitometric unit (RDU) analyses of β-Tubulin III protein levels were normalized by Coomassie blue (loading control) and corrected by control; UW228 (left panel) and D283 (right panel); *n* = 3 independent experiments. Data are shown as mean ± SEM; \**p* < 0.05, \*\**p* < 0.01, \*\*\**p* < 0.001, \*\*\*\**p* < 0.0001 compared to controls.

## DISCUSSION

We have previously reported (Thomaz et al., 2016) that ANA-12, a small molecule selective TrkB inhibitor (Cazorla et al., 2011), decreased cell viability, survival and promoted cell cycle arrest in MB cells *in vitro*. To increase our understanding of the functional role of TrkB in MB, here we evaluated cellular changes possibly associated with the antitumoral effects of TrkB inhibition *in vitro* and *in vivo*. ANA-12 has been successfully used to target TrkB in preclinical studies of other types of tumors, including lung cancer, Ewing sarcoma, glioblastoma, oral squamous carcinoma and leukemia (Polakowski et al., 2014; Sinkevicius et al., 2014; Heinen et al., 2016; Pinheiro et al., 2017; Moriwaki et al., 2018). The present study is the first to evaluate the antitumoral effects of ANA-12 in a pediatric cancer animal model. Overall, it shows that pharmacological TrkB inhibition by ANA-12 hinders the activation of TrkB downstream targets, such as ERK signaling, modulates gene expression, and leads to decreased MB cell viability and proliferation, induction of apoptosis, and promotion of morphological and molecular changes that are consistent with differentiation in MB cells. Two cell lines representative of different MB molecular subgroups were used in our study: UW228 cells are TP53-mutated and classified as Shh, whereas D283 cells are classified as Group 3/4 (Ivanov et al., 2016).

We found that MB cells treated with ANA-12 showed a decrease in the activation of ERK signaling pathway as measured by phospho-ERK. The ERK/MAPK cascade mediates the stimulatory effects of TrkB activation on proliferation (Chao, 2003; Thiele et al., 2009), and aberrant ERK signaling activation promotes proliferative stimuli in Shh and Group 4 MB (Badodi et al., 2017). ANA-12 is able to reduce ERK activation induced by BDNF in astrocytes (Saba et al., 2018) and prevent ERK activation in neurons (Brady et al., 2018). These findings are consistent with the possibility that blocking TrkB activity suppresses ERK-mediated growth and survival actions in MB.

STAT3 mRNA levels were increased by treatment with ANA-12, whereas phosphor-AKT levels remained unchanged. STAT3 plays an important role as an intermediate molecule linked to TrkB and AKT activation (Radin and Patel, 2017). MB tumors commonly present deregulation and aberrant expression of AKT and STAT3 and the activation of these pathways are associated with enhanced cellular survival, migration, angiogenesis and resistance to chemotherapeutic agents (Hartmann et al., 2006; Xiao et al., 2015). MB brain tumor-initiating cells expressing CD133 drive recurrence mediated by STAT3 activation (Garg et al., 2017). Thus, the upregulation in STAT3 after anticancer treatment with a TrkB inhibitor suggested by our findings could indicate a compensatory response to counteract the inhibitory effects. In fact, inhibition of receptor tyrosine kinases including the epidermal growth factor receptor (EGFR) and HER2 in experimental cancer models can trigger feedback activation of STAT3 as a possible mechanism of resistance to targeted therapies (Zhao et al., 2016). However, other studies indicate a dual role of STAT3, where it can act as a tumor suppressor. For instance, endogenous STAT3 activation or expression of a constitutively active form of STAT3 inhibit glioblastoma cell proliferation and invasiveness (de la Iglesia et al., 2008; Luwor et al., 2013). Thus, the increase in STAT3 expression we observed in ANA-12-treated cells may contribute to the antiproliferative effect of TrkB inhibition.

Inhibiting TrkB resulted in differential modulatory effects on p21 expression in the two MB cell lines used, where UW228 cells treated with ANA-12 showed an increase in p21 transcriptional expression whereas a decrease was found in D283 cells. It is possible that the increased expression of p21 in UW228 cells is associated with growth arrest, whereas decreased expression of p21 in D283 cells might be related to apoptosis. In addition, UW228 cells are TP53 mutated while D283 cells display TP53 expression (Ivanov et al., 2016). p21 displays a complex pattern of actions in regulating the cell cycle and interacting with other signaling components, and its role and expression may be different depending on p53 expression (Abbas and Dutta, 2009).

TrkB plays a role in cell differentiation during normal CNS development. We found that UW228 cells exposed to ANA-12 showed increased cellular complexity. Consistently with the observed morphological alterations, UW228 cells showed a small increase in mRNA levels of TUBB3 after TrkB inhibition. Upregulation of TUBB3 was previously associated with differentiation in MB cells (Fiaschetti et al., 2014). Moreover, a decreased expression of Nestin, a marker of stemness, was detected in both MB cells treated with ANA-12. Nestin is commonly used as a marker for neural stem cell and cancer stem cell populations (Neradil and Veselska, 2015), and its expression increases progressively during MB development (Li et al., 2013). Nestin cooperates with the Shh pathway to drive tumor growth and Nestin suppression inhibits cell proliferation and promotes differentiation in MB (Li et al., 2016). Thus, our findings provide early evidence that TrkB might be a target to induce differentiation in MB, particularly in Shh tumors, a possibility that should be further explored by future studies.

A recent study used ANA-12 as an experimental treatment for oral squamous carcinoma *in vivo* and found reduction of tumor growth in a pattern consistent with our results in MB (Moriwaki et al., 2018). Other TrkB inhibitors has also been shown to efficiently reduce the growth of neuroblastoma in mouse models (Croucher et al., 2015; Li et al., 2015). Our final series of experiments provides the first evidence that TrkB inhibition slows MB growth *in vivo*.

Given the critical role of BDNF/TrkB signalling in brain development and plasticity, concerns may arise regarding potential adverse effects of TrkB inhibitors on nervous system function in MB patients. Although this issue must be carefully considered and investigated, recent clinical studies of the pan-Trk inhibitor larotrectinib in children with NTRK gene fusion-positive solid tumors has suggested a good safety profile (Drilon et al., 2018; Laetsch et al., 2018).

Together, our results support the view that specific inhibition of TrkB can be effective as a therapy for MB, likely through mechanisms involving modulation of apoptosis and cell differentiation.

## AUTHOR CONTRIBUTIONS

AT, KP, BS, and MJ, performed the experiments. AT, ATB, VR, CN, and RR drafted the manuscript. All authors contributed to designing the experiments, analysing and interpreting the data, and revising and approving the final version of the manuscript.

## FUNDING

This research was supported by the National Council for Scientific and Technological Development (CNPq; grant numbers 303276/2013-4 and 409287/2016-4 to R.R. and grant number 201001/2014-4 to C.N.); PRONON/Ministry of Health, Brazil (number 25000.162.034/2014-21); the Children’s Cancer Institute (ICI); the Rio Grande do Sul State Research Foundation (FAPERGS; grant number 17/2551-0001 071-0 to R.R.); the Coordination for the Improvement of Higher Education Personnel; and the Clinical Hospital institutional research fund (FIPE/HCPA). C.N. is also supported by the William Donald Nash fellowship from the Brain Tumour Foundation of Canada. V.R. is supported by operating funds from the Canadian Institutes for Health Research, the American Brain Tumor Association, and the Brain Tumour Foundation of Canada. M.D.T. is supported by the National Institutes of Health, the Pediatric Brain Tumour Foundation, the Terry Fox Research Institute, the Canadian Institutes of Health Research, The Cure Search Foundation, b.r.a.i.n.child, Meagan’s Walk, Genome Canada, Genome BC, the Ontario Institute for Cancer Research, and the Canadian Cancer Society Research Institute.

## ACKNOWLEDGEMENTS

This manuscript has been released as a pre-print at bioRxiv: https://www.biorxiv.org/content/10.1101/566000v2.

