## Supplementary Material for "Antitumor Activities and Cellular Changes Induced by TrkB Inhibition in Medulloblastoma"

*[1]Cancer and Neurobiology Laboratory, Experimental Research Center, Clinical Hospital (CPE-HCPA), Federal University of Rio Grande do Sul, Porto Alegre, RS, Brazil, [2] Department of Pharmacology, Institute for Basic Health Sciences, Federal University of Rio Grande do Sul, Porto Alegre, RS, Brazil, [3] Children's Cancer Institute, Porto Alegre, RS, Brazil, [4] Department of Pediatrics, School of Medicine, Federal University of Rio Grande do Sul, Porto Alegre, RS, Brazil, [5] Pediatric Oncology Service, Clinical Hospital, Federal University of Rio Grande do Sul, Porto Alegre, RS, Brazil, [6] The Arthur and Sonia Labatt Brain Tumour Research Centre, The Hospital for Sick Children, Toronto, ON, Canada, [7] Division of Haematology/Oncology, The Hospital for Sick Children, Toronto, ON, Canada,[8] Developmental and Stem Cell Biology Program, The Hospital for Sick Children, Toronto, ON, Canada, [9] Department of Laboratory Medicine and Pathobiology, University of Toronto, Toronto, ON, Canada, [10] Division of Neurosurgery, The Hospital for Sick Children, Toronto, ON, Canada.*

***** Correspondence:***

*Rafael Roesler, Department of Pharmacology, Institute for Basic Health Sciences, Federal*

*University of Rio Grande do Sul, Rua Sarmento Leite, 500 (ICBS, Campus Centro/UFRGS), 90050-170 Porto Alegre,RS, Brazil.*

**

**SUPPLEMENTARY FIGURE S1** Original Western blots membranes analysed by ImageQuant LAS500 (GE Healthcare Life Sciences, Little Chalfont, UK) and original membranes stained with Coomassie blue (0.025%) for western blots in Figures 5 and 6. Membranes were cut prior to antibody stainings to allow for detection of proteins running at different sizes on the same membrane.

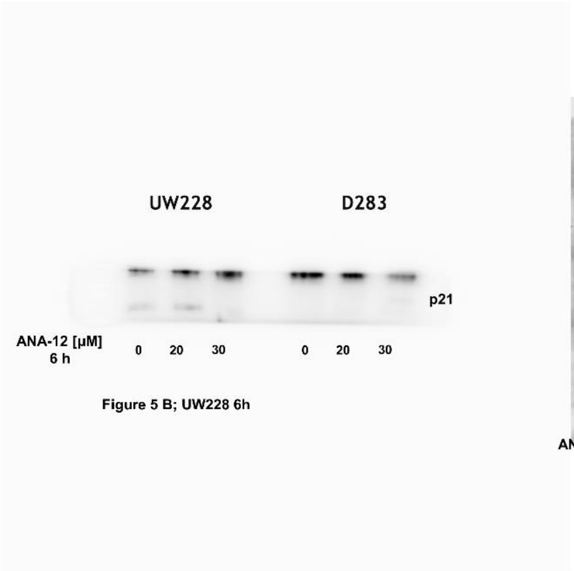


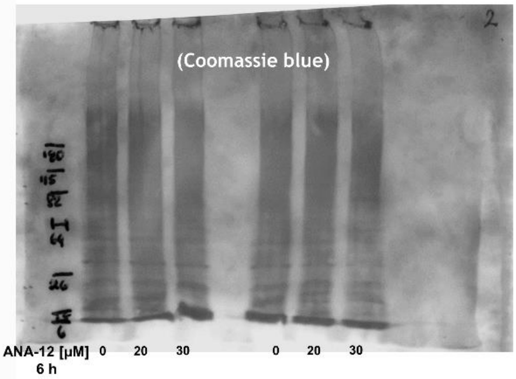


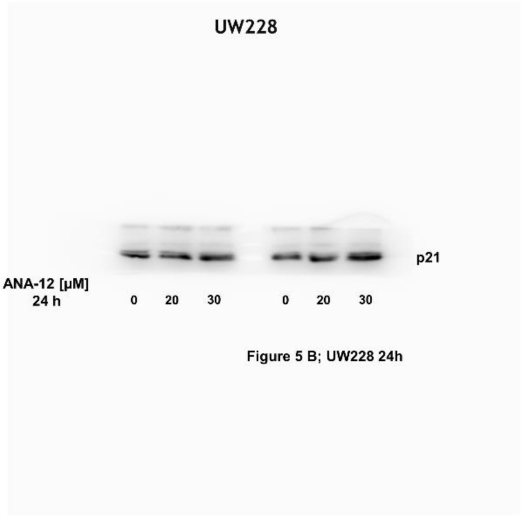


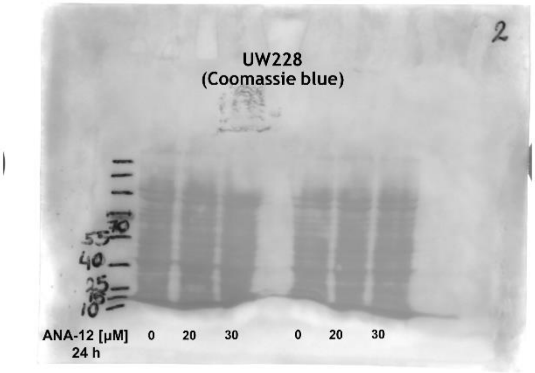

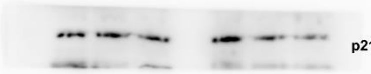


Figure 5 B; D283 6h

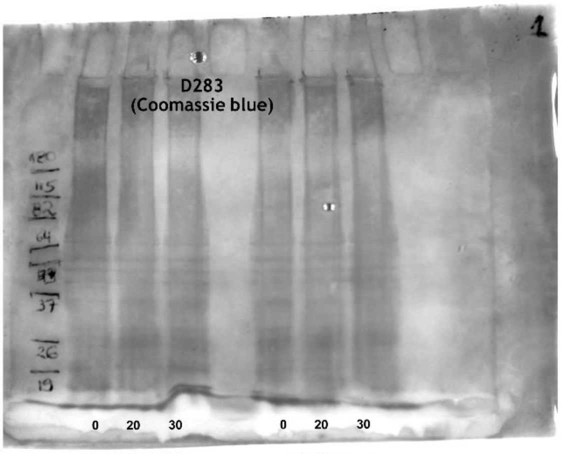


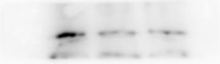


Figure 5 B; D283 24h

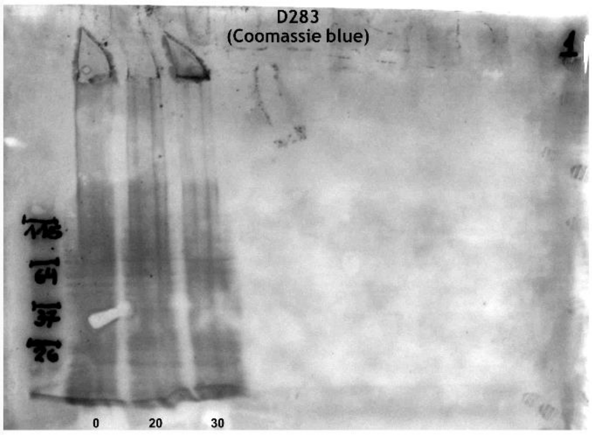

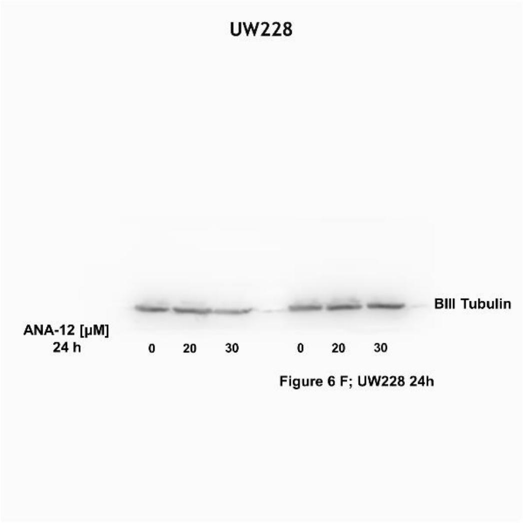

Figure 6 F; UW228 24h

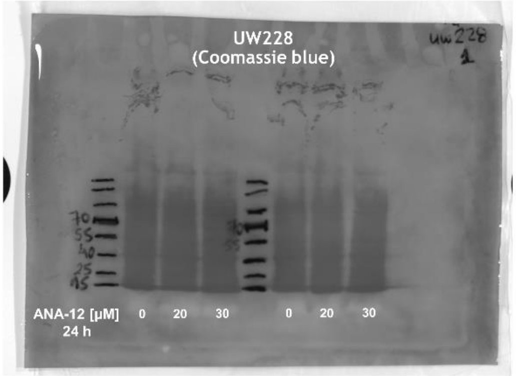


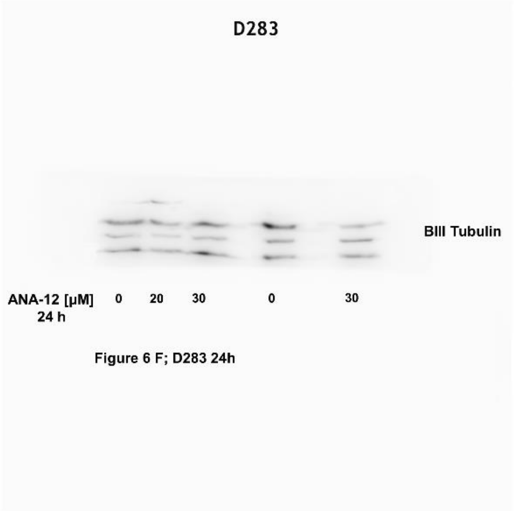

Figure 6 F; D283 24h

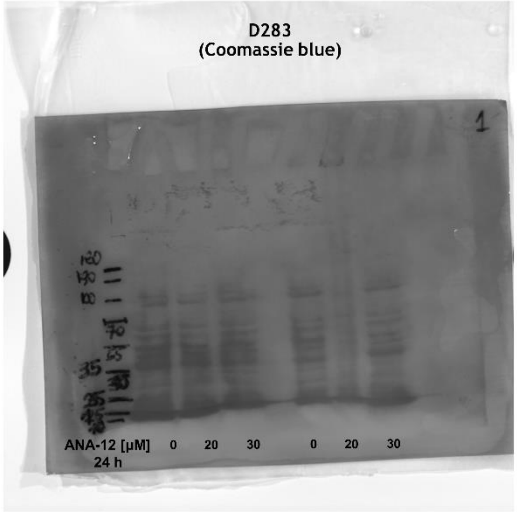
